# Noc1 Downregulation Induces Nucleolar Stress and Upregulates p53 Isoforms, with a Robust Increase of the Truncated p53E Isoform in Drosophila Wing Discs

**DOI:** 10.1101/2025.08.07.669174

**Authors:** Andrea Vutera Cuda, Shivani Bajaj, Valeria Manara, Paola Bellosta

**Affiliations:** Department of Cellular, Computational and Integrative Biology (CIBIO), University of Trento, Trento, TN 38123, Italy; Department of Medicine, NYU Langone Medical Center, New York, NY 10016, United States

**Keywords:** p53 isoforms, nucleolar stress, Noc1, apoptosis, γ-H2AV, primer specificity, isoform primer design, mutant screen, *Drosophila* imaginal discs

## Abstract

Disruption of ribosome biogenesis triggers nucleolar stress, a conserved cellular response that activates p53. We previously demonstrated that depletion of Nucleolar Complex Protein 1 (Noc1) in Drosophila wing imaginal discs impairs rRNA maturation and ribosome assembly, resulting in elevated p53 levels and apoptosis, hallmarks of nucleolar stress. The Drosophila p53 gene produces four mRNA isoforms, yet their individual contributions to nucleolar stress responses remain poorly understood. Using newly designed isoform-specific qPCR primers, we found that although all p53 isoforms exhibit moderate transcriptional changes following Noc1 reduction, the truncated isoform p53E is robustly and preferentially upregulated. Notably, p53E lacks the N-terminal transactivation domain and has been reported to negatively regulate p53-induced apoptosis in specific tissues. Furthermore, our analyses indicate that γ-H2AV accumulation arises from caspase-dependent apoptosis rather than primary genomic lesions, suggesting the activation of a p53-dependent stress pathway distinct from canonical genotoxic pathways. Together, these findings suggest that p53E may be part of a novel mechanism activated during nucleolar stress, providing insight into how cells adapt to defects in ribosome biogenesis.

## Introduction

Nucleolar stress is a cellular response to unbalanced ribosomal biogenesis (Boukoura and Larsen 2024). The nucleolus acts as a key stress sensor, responding to perturbations in ribosomal RNA (rRNA) processing and ribosome biogenesis by inducing the upregulation of p53 (Lindstrom et al. 2022).

*Drosophila* p53 exhibits significant structural and functional similarities with human p53 (Brodsky et al. 2000; Ingaramo et al. 2018; Ollmann et al. 2000; Bourdon et al. 2005). The *p53* gene (FBgn0039044) maps to chromosome 3R and encodes four mRNA isoforms (*p53A, p53B, p53C,* and *p53E*). According to the current deep RNA sequencing-based genome annotation, these isoforms encode three distinct proteins (Boley et al. 2014). Notably, *p53A,* also known as ΔNp53, and *p53C* differ in their 5’ untranslated regions (5’UTRs), and while the protein product of p53C remains unconfirmed, its open reading frame (ORF) is predicted to be identical to that of p53A, and to encode a 44 KDa protein (Zhang et al. 2015; Chakravarti et al. 2022). This suggests that differential promoter regulation underlies their distinct functions in different cellular contexts. *p53B* is the longer isoform, carrying a unique N-terminal domain and encoding for a 56 KDa protein considered the closest homolog to human p53 (TP53) (Ingaramo et al. 2018). *p53E* is the shorter isoform that encodes a predicted 38 kDa protein lacking part of the N-terminal transactivation domain, and is considered to act as a dominant negative in specific tissues and stress conditions (Wylie et al. 2022).

p53A, and p53B are the most characterized protein isoforms. While their expression is comparable in the germline, they begin to exhibit distinct expression and regulation patterns across various developmental stages (Chakravarti et al. 2022). In somatic tissues, p53A mRNA and protein were found to be more abundant, likely reflecting differential roles during tissue development (Chakravarti et al. 2022). This may reflect their role in controlling apoptosis.

Initial studies demonstrated that overexpression of each p53 protein isoform leads to embryonic lethality. In addition, it was demonstrated that overexpression of p53A or p53B and of p53E induced apoptosis in the developing eye when driven by the GMR promoter, while in the ovary, overexpression of p53A and p53B proteins markedly enhanced apoptosis following irradiation, whereas p53E overexpression reduced the number of apoptotic cells, suggesting a dominant-negative role for this isoform in the ovary under irradiation-induced stress conditions (Zhang et al. 2015). Numerous studies have demonstrated that both p53 protein A and B isoforms play distinct roles in regulating stress responses and inducing apoptosis; indeed, p53A was shown to be both necessary and sufficient to activate the proapoptotic genes *rpr* and *hid* in response to DNA damage in somatic and germline tissues (Zhang et al. 2015; Chakravarti et al. 2022).

Although p53B can induce apoptosis when experimentally overexpressed, its behavior in normal tissues appears to be different. In several somatic tissues, the mRNA levels of p53B are only slightly lower than those of p53A, yet the amount of p53B protein detected is considerably lower. This discrepancy suggests that p53B may be translated less efficiently or undergo more rapid protein degradation (Zhang et al. 2015; Zhang et al. 2014). These differences in protein abundance and activity among tissues may stem from competition between p53 isoforms that have opposing regulatory functions. Specifically, p53B primarily acts as a transactivator, promoting the expression of target genes, whereas p53A, which is more broadly expressed across tissues, can function as both a transactivator and a transrepressor, (Wylie et al. 2022). Thus, the relative levels and functional properties of these isoforms likely contribute to the tissue-specific outcomes of p53 signaling. In support of this hypothesis, studies in photoreceptor neurons have shown that the balance between autophagy-mediated survival and apoptosis-driven cell death is governed by antagonistic regulatory inputs from p53A and p53B, whose mRNA levels shift differentially in response to oxidative stress (Robin et al. 2019).

Although p53 is best known for its conserved role in apoptosis triggered by DNA damage, it also exerts broad control over tissue physiology and metabolic homeostasis. Under nutrient-limiting conditions, *p53*-null mutant flies fail to sustain normal survival, revealing that p53 is essential for coordinating metabolic adaptation and efficient nutrient consumption during stress (Barrio et al. 2014). p53 also regulates growth and proliferation in a coordinated, non-autonomous manner and modulates apoptosis in response to stress, such as rRNA depletion (Mesquita et al. 2010). Furthermore, p53 is necessary to maintain imaginal disc plasticity and regenerative potential (Wells and Johnston 2012) and to act as a general sensor of competitive confrontation during MYC-induced cell competition (de la Cova et al. 2014). Notably, p53 isoform-specific protein functions have not been assessed in these experiments despite growing evidence that their function can differ significantly depending on the cellular context in which they are expressed. Thus, it may be important to consider that distinct signals and tissue-specific expression programs could drive isoform-specific roles in stress response and gene regulation.

Our previous study demonstrates that reducing the nucleolar gene *Noc1* (FBgn0036124) impairs rRNA processing and ribosome biogenesis, leading to apoptosis and the upregulation of p53 (Rambaldelli et al. 2025; Destefanis et al. 2022), both hallmarks of nucleolar stress. Using our newly designed p53 isoform-specific qPCR primers, we found that *p53E* is the most upregulated isoform following *Noc1* reduction. This suggests that nucleolar stress may induce a novel regulatory mechanism for p53E, whose strong upregulation upon nucleolar disruption points to a potential autoregulatory interaction in which one p53 isoform may influence the activity or stability of other isoforms. In addition, γ-H2AV phosphorylation accumulates because of caspase-dependent cell death rather than direct genomic lesions, consistent with the activation of a p53 stress pathway distinct from the canonical genotoxic pathway. Together, these findings suggest that p53E may be part of a novel mechanism activated during nucleolar stress, providing insight into how cells adapt to defects in ribosome biogenesis.

## Materials and Methods

### Drosophila husbandry and lines

Animals were raised at low density in vials containing standard fly food, composed of 9 g/l agar, 75 g/l corn flour, 50 g/l fresh yeast, 30 g/l yeast extract, 50 g/l white sugar, and 30 ml/l molasses, along with nipagin (in ethanol) and propionic acid. The crosses and flies used for the experiments are kept at 25°C, unless otherwise stated.

Line used: *w^1118^* from VDRC #6000, *Rotund*-Gal4 BDSC#76179, *UAS-Noc1-RNAi* BDSC #25992, *UAS-p35* (Destefanis et al. 2022), *p53^5-1-4^* BDSC #6815, *p53^A2.3^* and *p53^B41.5^* from (Chakravarti et al. 2022).

### Alkaline Comet Assay

Using the *UAS-Gal4* system (Brand and Perrimon 1993), *Noc1* was downregulated in the entire imaginal disc using a temporally controlled expression system under the *Actin-Gal4* promoter in combination with the temperature-sensitive *Gal80-ts* allele that inhibits Gal4 (Germani et al. 2018). Animals were left to develop for five days after egg laying (AEL) at the permissive temperature of 18°C, then shifted to 30°C for 60 hours, during which Gal4 was active and induced the ubiquitous expression of *Noc1-RNAi*. Wing imaginal discs were dissected and incubated in Ringer’s solution, then enzymatically digested with Collagenase in TrypLE at 37–38 °C for 30 minutes following the protocol in (Pederzolli et al. 2025). After electrophoresis and Hoechst staining, photos of nuclei were imaged using a Zeiss Axio Imager fluorescent microscope with a 40X objective, and DNA damage was analyzed with CometScore 2.0.

### RNA Extraction and RT-qPCR

RNA extraction was performed with the RNeasy Mini Kit (Qiagen). Five larvae or 30 wing discs were collected and placed in 250µl of lysis buffer, then processed following the manufacturer’s instructions. The isolated RNA was quantified using a Nanodrop (Thermo Scientific) and stored at −80 °C for extended periods. RNA (1µg) was retro-transcribed into cDNA using the SuperScript IV VILO Master Mix (Invitrogen). DNase was added to the samples. To perform retro-transcription, the mix was incubated at 25°C for 10 minutes, 50°C for 10 minutes, and 85°C for 5 minutes. The obtained cDNA was used to perform quantitative PCR (qPCR) with the SYBR Green PCR Kit (Qiagen). The final mix for each sample to be tested was composed by 8µl of cDNA, 2µl of primers (10µM), and 10µl of SYBR Green reaction mix. Reactions were carried out using a BioRad CFX96 machine with the following protocol: 50°C for 2 minutes, 95°C for 2 minutes, 95°C for 15 seconds, and 60°C for 30 seconds (repeated for 45 cycles), followed by 72°C for 1 minute. Analyses were done using Bio-Rad CFX Manager software. The relative abundance level for each transcript was calculated by subtracting the value of actin from the value of p53 and using the 2^-ΔΔCt^ method. For qPCR in wing discs analysis, the RNA was extracted from at least 20 wing imaginal discs, or from at least five larvae for each genotype. Data are expressed as arbitrary units (A.U.). The number of independent biological replicates is cited in the figure legend for each data set. The sequences of the primers used are in Table 1.

**Table 1:**
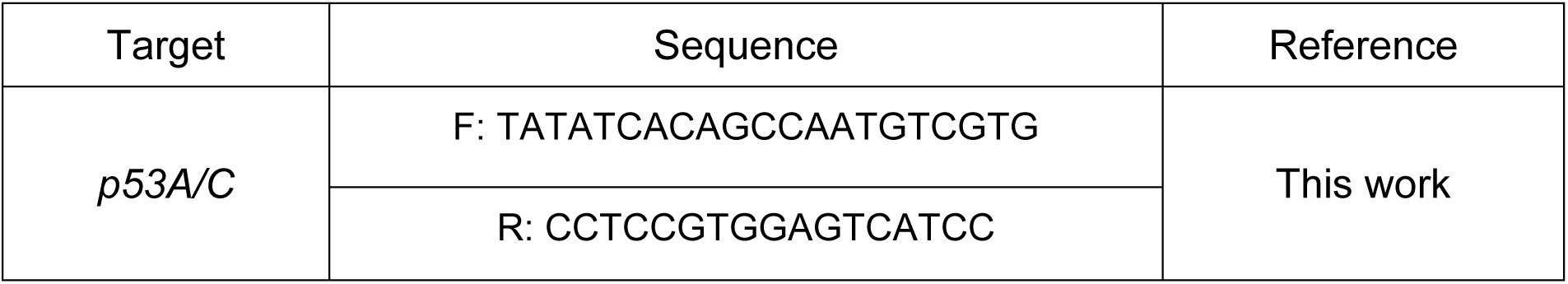
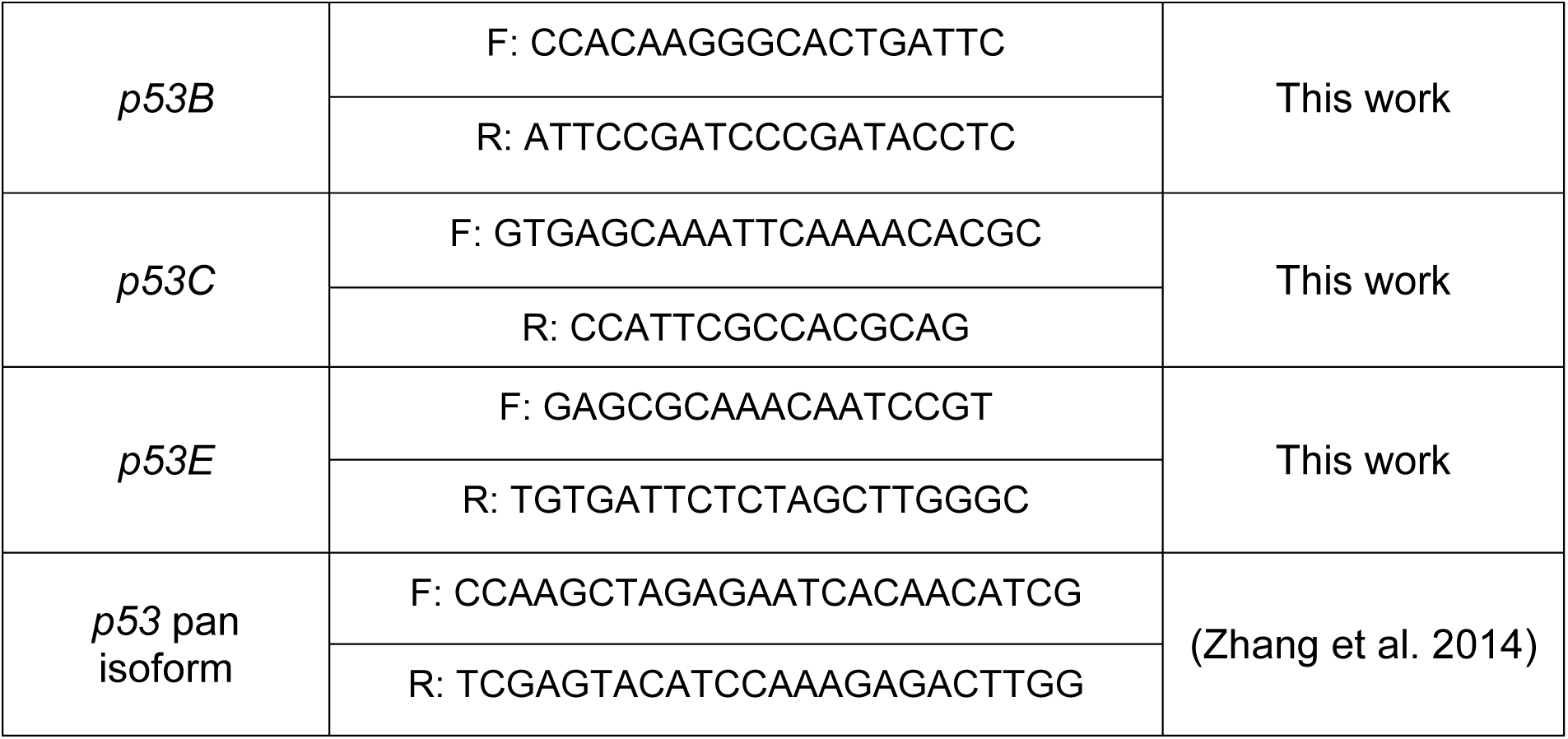
Sequences of the primers used in this work.

### Dissections and immunofluorescence

Larvae were collected at the third-instar stage, dissected in 1× phosphate-buffered saline (PBS), and fixed for 30 min in 4% paraformaldehyde (PFA) at room temperature (RT). After 15 minutes of tissue permeabilization with 0.3% Triton X-100, samples were washed in PBS with 0.04% Tween 20 (PBS-T) and blocked in 1% bovine serum albumin (BSA) for 1 hour at RT. Samples were incubated overnight at 4°C with primary antibodies in 1% BSA and, after washing, with Alexa-Fluor-488- or 555-conjugated secondary antibodies in BSA. During washing in PBS-T, nuclei were stained with Hoechst 33342 (Thermo Fisher Scientific). Imaginal discs were dissected from the carcasses and mounted on slides with Vectashield (LsBio-Vector Laboratories). Images were acquired using a Leica SP8 confocal microscope or Zeiss Axio Imager fluorescent microscope and analyzed with the Fiji program to measure fluorescence. Images were assembled using Adobe Photoshop 2025. Primary antibodies used were rabbit anti-cleaved Caspase3, 1:400 (Cell Signaling 9661); and anti p2HAV, 1:1000 (DSHB#UNC93-5.2.1)(Lake et al. 2013).

### Image processing

Raw confocal images were processed with Fiji to obtain a z-stack projection (using the Average Intensity or Max Intensity options). Then, all images from the same experiment were processed, enhancing the brightness of all channels equally across all images.

## Results

### 1.1 Reduction of Noc1 triggers DNA Double-Strand Breaks, p53 upregulation, and apoptotic signaling that regulates γH2AV phosphorylation

We previously demonstrated that downregulation of the nucleolar factor Noc1 in *Drosophila* tissues leads to the accumulation of rRNA (Destefanis et al. 2022; Rambaldelli et al. 2025). Furthermore, *Noc1-RNAi* expression driven by the *rotund* promoter in cells of the wing imaginal discs induces DNA damage as evidenced by the comet-like morphology of the nuclei in the affected cells (Figure 1A-B, and (Pederzolli et al. 2025). These nuclei show a statistically significant increase in the comet tail length (or tail moment, calculated as tail length multiplied by the percentage of DNA in the comet head), indicating the presence of double-strand breaks absent in the nuclei of wild-type *w^1118^* cells (Figure 1C). Notably, reduction of Noc1 is accompanied by a significant increase in apoptosis in cells of the wing pouch (Figure 1D-E and F), together with the upregulation of p53 at both transcriptional and protein levels (Figure 1G-H).

**Figure 1.**
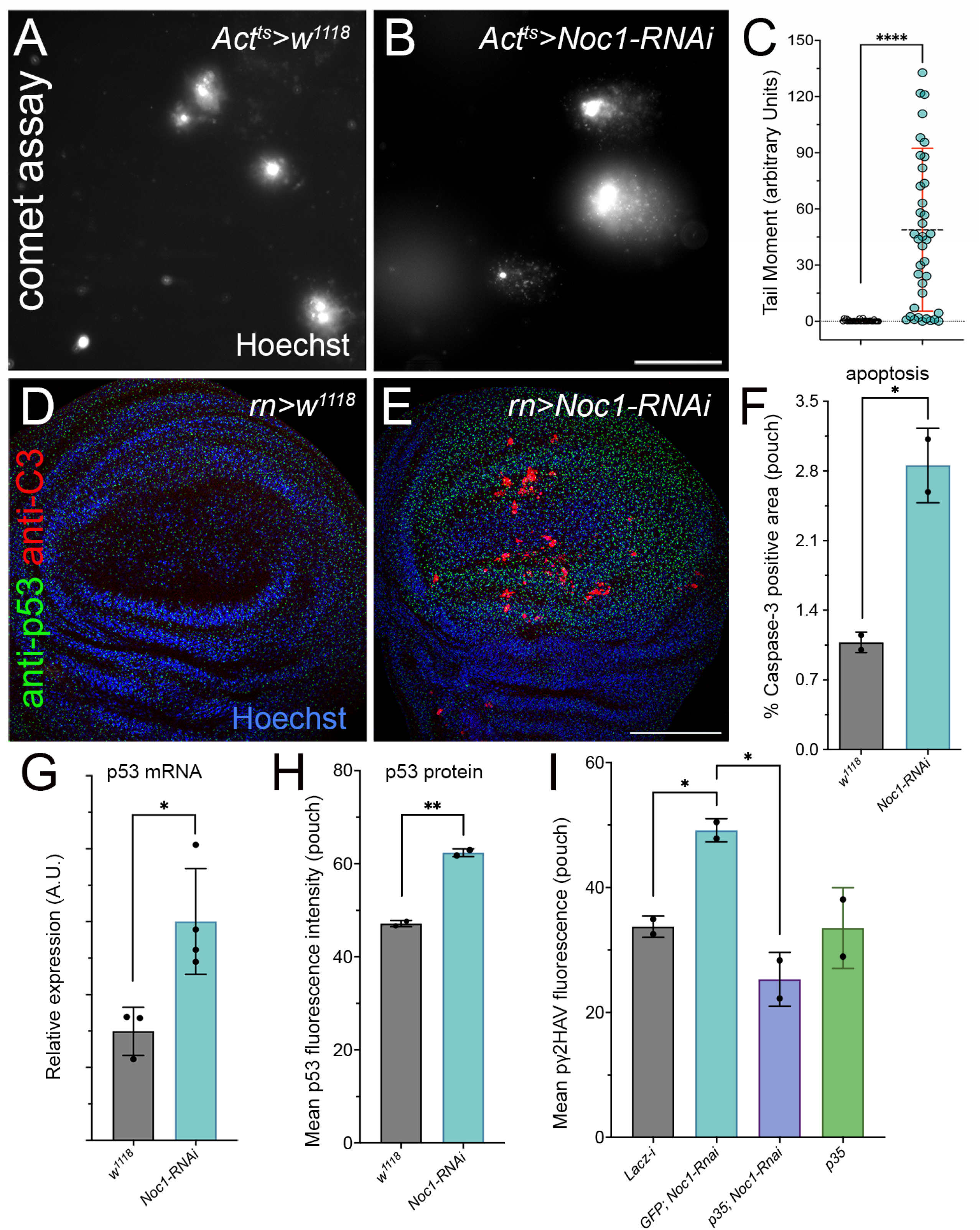
Noc1 Downregulation induces DNA double-strand breaks (DSBs), leading to p53 upregulation and apoptosis, which contributes to γH2AV phosphorylation. (A-C) Comet assay showing fluorescent images of nuclei stained with Hoechst (white) from cells of the wing imaginal discs of *w^1118^* third instar larvae (A), and from animals in which *Noc1* was transiently reduced for 60 hours under the temporal control of the *Act-Gal80^ts^* system. (B), scale bar is 50*μ*m. (C) Quantification of single-cell tail moment from the comet assay performed on nuclei from the genotypes shown in A and B. These data are obtained from pooling at least 20 wing imaginal discs of each genotype, and one representative experiment is shown. Statistical analysis was performed using the non-parametric Kruskal-Wallis as described in (Pederzolli et al. 2025). Experiments in (D-I) are from wing imaginal discs from third instar control larvae (*rn>w^1118^*) or from larvae in which expression of the transgenes was reduced using the indicated Rnai lines in the wing pouch under the control of the *rotund* (*rn*) promoter. (D-E) Confocal images of wing imaginal discs from third instar larvae immunostained for the expression of p53 protein (green) and for Caspase3, as a marker of apoptosis (red). Nuclei were stained with Hoechst (blue). Scale bar is 100*μ*m. (F) Quantification of Caspase3 by immunofluorescence in cells from the pouch of wing discs from third instar larvae of the indicated genotype. (G) qPCR analysis from RNA extracted from at least 20 wing imaginal discs from larvae of the indicated genotype, *p53* mRNA expression was analyzed using p53 pan isoform primers (Table 1). Data are expressed as arbitrary units (A.U.) (H). Quantification of p53 by immunofluorescence was analyzed in the pouch of wing imaginal discs from animals of the indicated genotype. (I) Quantification of γH2AV phosphorylation by immunofluorescence was analyzed in the wing pouch form larvae of the indicated genotype. In C, the asterisks represent the *p-values* from Student’s *t*-test **** = *p* < 0.0001, in one out of three independent experiments performed, where the dots represent the number of nuclei analyzed. In F, G, H, and I, the dots represent the number of independent biological replicates, with at least five imaginal discs analyzed for each genotype. The asterisks represent the *p-values* from Student’s *t*-test * = *p* < 0.05 and ** = *p* < 0.01; the error bars indicate the standard deviations.

To assess the impact of Noc1 downregulation on DNA damage signaling, we monitored γH2Av phosphorylation, a well-established marker of double-strand breaks. Interestingly, blocking apoptosis prevented γH2AV accumulation (Figure 1I), indicating that in this context, γH2Av phosphorylation is downstream of apoptotic signaling rather than a direct consequence of Noc1 loss. This suggests that apoptosis itself contributes to the observed DNA damage response in these tissues. Together, these findings are consistent with a cellular response to nucleolar stress resulting from impaired rRNA processing upon Noc1 reduction. Based on these data, we designed isoform-specific qPCR primers to determine which individual p53 isoforms respond to Noc1 reduction, as they may respond specifically to nucleolar stress.

### 1.2 p53 locus and novel primers for detecting distinct p53 isoforms

The *Drosophila* genome contains a single *p53* locus (Figure 2A, FBgn0039044 and NCBI gene ID: 2768677) on chromosome 3R, which produces four distinct mRNA transcripts through alternative promoter usage and splicing, resulting in four protein isoforms (Figure 2B). The canonical *p53A* (also referred to as DNp53) lacks part of the N-terminal Transactivation Domain (TAD), producing a protein of approximately 44 kDa. The *p53B* transcript retains the full TAD, encoding for a protein of about 56 kDa, while the rest of the sequence is identical to that of *p53A* mRNA. p53C protein shares the same amino acid sequence as p53A, but their mRNAs originate from an alternative transcription start site and are predicted to produce a longer transcript. Finally, *p53E* is the shortest isoform, predicted to encode a 38 kDa protein that lacks the N-terminal TADs of p53A and p53B, but retains the DNA-binding domain. This isoform has been shown to inhibit the apoptotic response to ionizing radiation, suggesting that it may act as a dominant-negative regulator of the other p53 isoforms (Wylie et al. 2022; Zhang et al. 2015).

**Figure 2.**
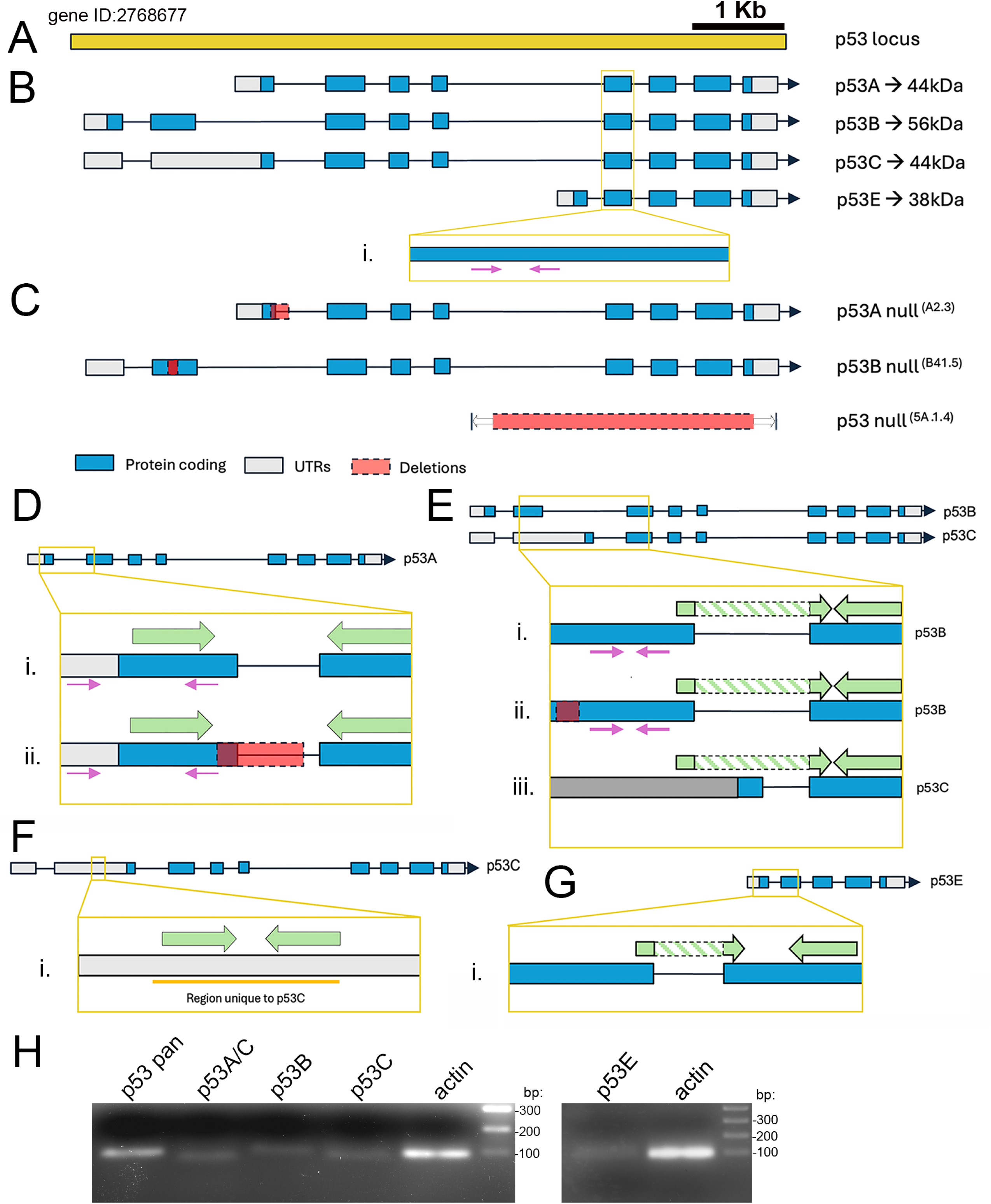
Schematic representation of the p53 locus and primer design used to detect specific isoforms. (A-B) Schematic representation of the *p53* locus (gene ID 2768677) and of the p53 isoforms *A, B, C,* and *E*. (B-i), magnification of a common exon with the representation of the region where the primers that capture all *p53* isoforms anneal (Zhang et al. 2015). These primers detect mRNA transcripts for all isoforms and were used in Figure 1F; they will be referred to as p53 pan. (C) Schematic representation of the *p53^A2.3^*, *p53^B41.5^,* and *p53^5A-1-4^* null mutant alleles. The *p53^A2.3^* null line was generated with a 23bp deletion and a 7bp insertion spanning from the end of the first exon until the first base of the first intron (in red), generating multiple early STOP codons (Supplementary Figure 3A). This disrupts protein production from the p53A and p53C isoforms, but mRNA may still be detectable if primers anneal outside the deleted region. *p53^B41.5^* null line carries a 14bp deletion and a 1bp insertion in exon 2, resulting in a frameshift and premature stop codons leading to a truncated, non-functional p53B protein (Supplementary Figure 3B)(Chakravarti et al. 2022). *p53^5A-1-4^* null line, contains a 3.3 kb deletion that removes the common 3’ coding region (in red), eliminating protein expression from all isoforms. (D) *p53A* isoform. (D-i and ii). Magnification of exons 1 and 2 of *p53A* and *p53^A2.3^* null isoform, respectively, showing where the primers designed in this study anneal (green) compared to previous primers (in pink). As shown in D-ii, the new primers can also be used to detect *p53A* mRNA in the *p53^A2.3^* null mutant, because the forward primer anneals outside the deleted region. In contrast, the previous reverse primer (pink) has 1 bp mismatch and may fail to detect the transcript in the *p53A* null line. The new primers also detect *p53C* mRNA and will therefore be referred to as p53A/C. (E) *p53B* and *p53C* isoforms. (E-i, ii, and iii). Magnification of exons 2 and 3, shared by both isoforms, showing where the primers designed in this study anneal (in green). The new primers specifically detect *p53B* mRNA, whereas the old primers (pink) also detect *p53C* mRNA. Moreover, the new pair is also suitable for analyzing the *p53B* null line because the forward primer does not anneal in the deleted region marked in red (E-ii). This pair of primers will be referred to as p53B. (F) *p53C* isoform. (F-i). Magnification of the region in exon 2 that includes the sequence of 78 bp, from 440 to 518 nucleotides from the beginning of the *p53C* transcript (in orange), unique to this isoform. Optimized primers (green) detect *p53C* mRNA and are referred to as p53C. (G) *p53E* isoform. (G-i). Magnification of the region between exons, highlighting the amplified region specific to this isoform. (H) Agarose gel (2%) showing qPCR-amplified products from RNA extracted from wing imaginal discs of *w^1118^* third-instar larvae, using the indicated primers and *actin* as a control. A single band is observed for each primer pair, demonstrating the specificity for mRNA detection.

Although primer sets have previously been reported for distinguishing among p53 mRNA isoforms (Supplementary Figure 1), none have been fully optimized to reliably differentiate between *p53A* and *p53B*. Additionally, primers capable of specifically discriminating between *p53B* and *p53C,* or uniquely amplifying *p53C*, remain unavailable. To overcome these limitations, we designed and optimized a novel set of primers to enable precise and isoform-specific analysis of p53 transcripts (Table 1).

To ensure isoform specificity and accurate validation, primers were designed using the sequences of the available *p53^5A-1-4^* null lines and the isoform-specific *p53A* and *p53B* mRNA mutant lines. Briefly, the *p53A* null line carries a 23bp deletion and 7bp insertion at the end of the first exon which expands also in the first intron (Robin et al. 2019); the mutant *p53B* null line has a 14 bp deletion and 1 bp insertion in the second exon, resulting in multiple early STOP codons (Chakravarti et al. 2022); and the *p53^5A-1-4^* null line carries a large deletion of approximately 3.3 Kb that removes four exons, preventing the transcription of a functioning mRNA, resulting in a complete loss of p53 protein expression (Figure 2C) (Rong et al. 2002).

The new primers for the *p53A* isoform span the exon-exon junction between exons 1 and 2 (Figure 2D-i, shown as light green arrows in the panel, and Table 1), providing enhanced specificity compared to earlier designs (shown as arrows in pink, and Table 1). Like the previously available primers, this set can be used to assess *p53A* expression in the p53A null line as the forward primer anneals upstream of the deleted region, while the reverse primer anneals right after the deletion, allowing the amplification of a specific region of the mutant (Figure 2D-ii). A limitation of the new primers, as with previously published ones, is their inability to distinguish between *p53A* and *p53C* transcripts, due to the high sequence similarity between them. To design primers capable of distinguishing *p53B* from *p53C* isoform, we took advantage of the difference in the length of their 5′ untranslated region (5’UTR) (Figure 2E). While the reverse primer anneals in a sequence common between *p53B* and *p53C*, the forward primer for *p53B* anneals at the end of the second exon and extends into the beginning of exon 3 (green arrows in Figure 2E-i and 2E-iii). This design maximizes specificity, ensuring amplification of only *p53B*. Moreover, this set of primers is suitable for screening *p53B* in *p53B* null lines, as they do not anneal within the deleted region (Figure 2E-ii).

Through detailed analysis of the 5′UTR sequence of *p53C*, we identified a region unique to this isoform (highlighted in orange in Figure 2Fi). Thus, we designed primers that specifically target this region, enabling the selective detection of *p53C* mRNA and providing a distinct primer set for this isoform. Furthermore, the availability of p53C-specific primers allows for a more accurate interpretation of results obtained with the p53A primer set, which cannot distinguish between the *p53A* and *p53C* transcripts.

In addition to p53A, p53B, and p53C, we developed isoform-specific primers for the truncated p53E isoform. These primers target the unique 5’ sequence generated by the p53E transcription start site (Figure 2G), providing selective amplification and preventing cross-detection with other isoforms.

Primer specificity was tested by performing qPCRs on RNA extracted from imaginal discs of *w^1118^* third-instar larvae. All primers amplified a single product, as evidenced by a single band observed in the gel (Figure 2H) and confirmed by melting curve analysis, which showed a distinct peak for each primer (Supplementary Figure 2).

### 1.3 Validation of isoform-specific primers in wild-type and p53 mutant background

To further assess the specificity and to complete the characterization of the newly designed primer sets, we conducted qPCR analysis on RNA from wild-type control *w^1118^*and *p53* mutant lines using primers targeting the single *p53* mRNA isoforms (Table 1).

As shown in Figure 3A, the p53 pan primers, which target a region common to all the *p53* isoforms (Figure 2B-i), detect p53 transcript levels in the *p53A* null line that are comparable to those of control *w^1118^*. In contrast, the *p53B* null line exhibits a small but not significant increase in *p53* mRNA expression, suggesting a potential compensatory upregulation in response to the loss of *p53B*.

**Figure 3.**
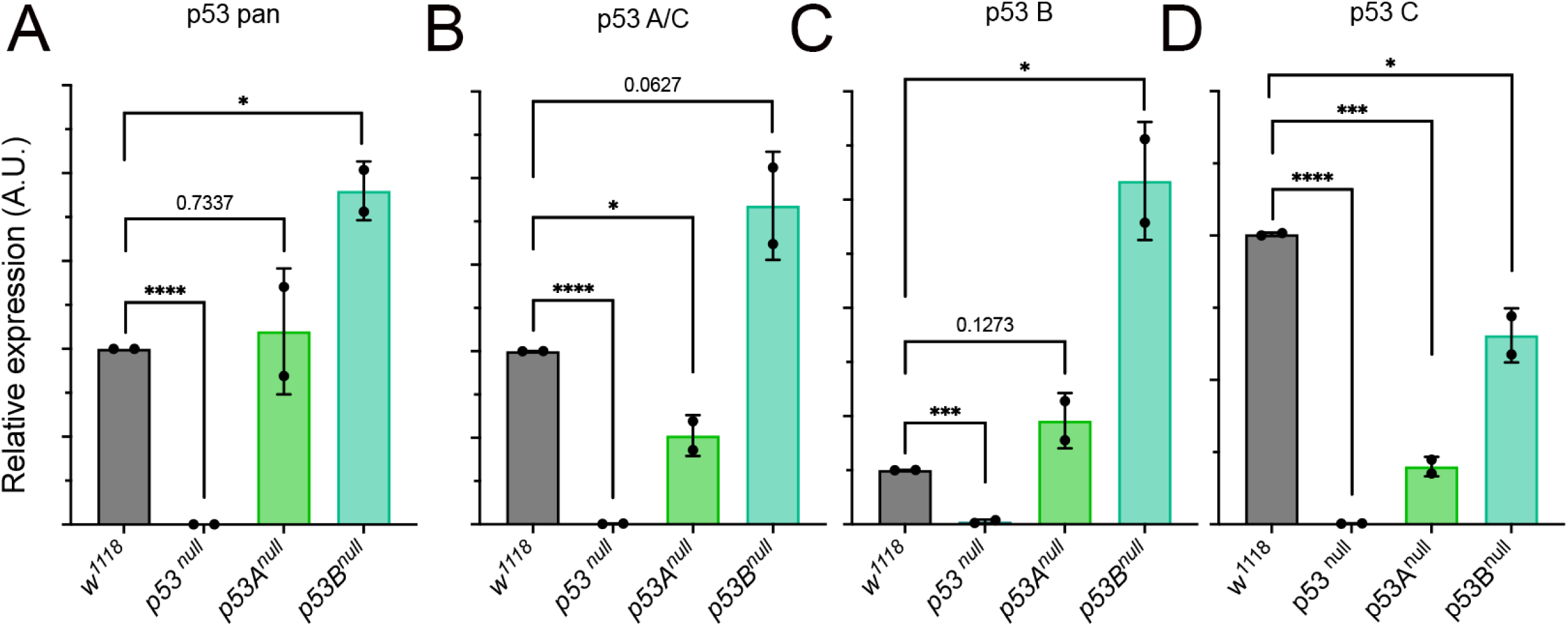
Expression analysis of *p53 mRNA* isoforms by RT-qPCR in p53 mutant lines using isoform-specific primers. (A-D) Relative expression of *p53 mRNA* isoforms compared to *actin5C,* used as a control. The relative p53 isoforms analyzed with the specific primers are indicated at the top of each graph. The genotype of the lines is shown below. Relative expression of *p53 mRNA* is shown as arbitrary units (A.U.). *p53* pan detects all p53 isoforms; p53A/C specifically detects *p53A* and the predicted *p53C* isoforms; p53B, p53D, and p53E detect only the respective isoform; *w^1118^* animals are used as control. The asterisks indicate the *p-*values obtained from Student’s t-test analysis from two independent biological experiments. RNA was extracted from five whole larvae of the respective genotype for each experiment. * = *p* < 0.05, *** = *p* < 0.001, and **** = *p* < 0.0001, and the error bars indicate the standard deviations.

Using the p53A/C primers, we observed that the *p53A/C* mRNA levels were significantly reduced by approximately 50% in the *p53A* null line (Figure 3B). In contrast, the *p53B* null line showed a significant twofold increase in *p53A/C* mRNA expression. Consistent with the data in Figure 3A, these findings may suggest the presence of a compensatory response to the other isoforms when p53B is eliminated in null animals, whereas loss of p53A has minimal effect on overall transcript abundance.

In the *p53B* null line (Figure 3C), we observed an unexpected significant upregulation of *p53B* mRNA. However, since the mutation does not prevent transcription and the primers amplify regions downstream of the deletion, it is likely that the mutant transcript can still be detected by qPCR. Overall, this finding suggests that compensatory or autoregulatory feedback mechanisms may act to increase *p53B* transcription or stabilize the mutant mRNA in response to reduced levels of p53B protein.

Finally, analysis of the *p53C* isoform, using primers that specifically detect the transcript, reveals a significant reduction in its expression in both *p53A* and *p53B* null lines (Figure 3D), suggesting that both *p53A* and *p53B* isoforms, directly or indirectly, participate in the regulation of *p53C* expression.

Importantly, the complete absence of *p53* transcripts in the *p53^5A-1-4^* null line confirms the specificity and reliability of the isoform-targeted primer set used in this study.

### 1.4 p53E is the predominantly upregulated isoform upon Noc1 downregulation

Analysis of p53 expression in cells of the wing imaginal disc in which Noc1 was reduced in the pouch revealed a significant upregulation of *p53* mRNA levels when assessed using pan-isoform primers (Figure 4A and Figure 1G). Because these primers recognize a conserved region shared by all four p53 isoforms, as illustrated in the magnified view (Figure 2B-i), this initial analysis did not allow discrimination between individual isoform contributions. Given that Noc1 reduction leads to rRNA accumulation and increased p53 levels (Figure 1B-C), and considering that p53 isoforms can be expressed at different levels under various stress conditions, we therefore designed isoform-specific primers to determine which p53 isoform(s) are preferentially upregulated in response to *Noc1-RNAi-*induced nucleolar stress.

**Figure 4.**
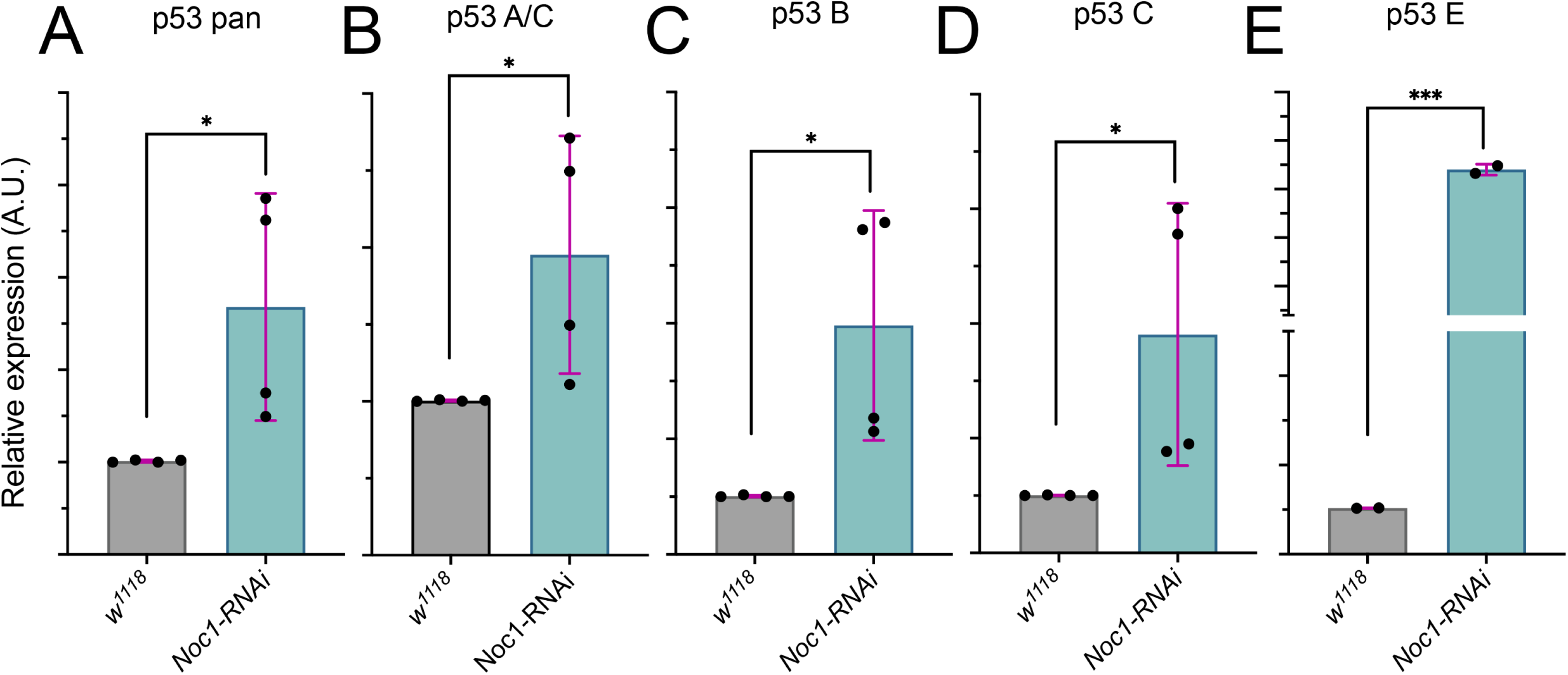
Upregulation of p53 isoforms in Noc1-downregulated cells. (A-E) qPCRs from RNA extracted from wing imaginal discs of *w^1118^* animals and expressing *Noc1-RNAi* in the wing pouch using the *rotund* promoter. Data are expressed as relative expression of *p53 mRNA* isoforms as arbitrary units (A.U.) compared to *actin5C,* used as a control. The asterisks represent the *p-*values obtained from Student’s t-test analysis: * = *p* < 0.05, and *** = *p* < 0.001. The error bars indicate the standard deviations. The dots represent the number of independent biological experiments. In each biological experiment, 30-40 wing imaginal discs were dissected per genotype and pooled for RNA extraction.

Among the isoforms, p53E exhibited the strongest induction (*p*< 0.001), whereas all the other isoforms showed a moderately significant increase (*p*< 0.05). Notably, p53E has been reported to act as a dominant negative in a tissue-specific manner. Thus, our analysis suggests that p53E may contribute to the repression of specific target genes during the nucleolar stress response.

Collectively, these findings highlight an isoform-specific regulation of p53 in response to nucleolar stress, with p53E emerging as a potential novel key mediator of transcriptional repression. The differential upregulation of p53 isoforms suggests a fine-tuned cellular strategy, where transrepressor isoforms may selectively modulate gene expression to orchestrate an appropriate stress response, while other isoforms potentially maintain canonical p53 functions. This underscores the importance of considering isoform-specific roles in understanding p53-mediated pathways under nucleolar stress conditions.

## Discussion

In this study, we present an optimized and validated set of isoform-specific primers designed for the accurate detection and quantification of *Drosophila* p53 transcript variants. Here, we also designed primers to specifically detect the unique p53C isoform, enabling precise isoform discrimination. We first utilized p53 mutant lines to validate the specificity of these primers. These analyses show that the *p53A* null line retains some of *p53A* and *p53C* transcript levels (Figure 3B and D). Moreover, despite the introduction of premature stop codons within the coding sequence of the *p53B* mutation, our qPCR analysis revealed an upregulation of *p53B-*mRNA in *p53A/C* and *B* mutant lines (Figure 3D and C), which may suggest the presence of a compensatory mechanism, to buffer the loss of p53B function, or to compensate for the reduction of p53A/C protein, highlighting isoform-specific plasticity of the p53 network, where alternative isoforms may adjust expression to preserve critical stress response functions.

Our analysis of isoform-specific p53 expression under nucleolar stress induced by Noc1 reduction (Milkereit et al. 2001; Destefanis et al. 2022) (Figure 1) reveals that all *p53* isoforms were upregulated (*p*< 0.05), with *p53E* showing the most robust increase (*p*< 0.01), marking its first correlation with nucleolar stress and its role in stress-induced apoptosis. This truncated variant, characterized by a unique N-terminal domain, has been shown to counteract p53-induced apoptosis in eye imaginal discs (Zhang et al. 2015).

Indeed, the transcriptional repression activity of p53 isoforms is highly context-dependent. While p53A was shown to act as both an activator and a repressor in salivary glands, p53E functions only as a transrepressor and is dispensable for transactivation in irradiated embryos, where p53A was essential (Wylie et al. 2022). However, our study specifically addresses nucleolar stress, which is mechanistically distinct from stress induced by DNA damage. In wing imaginal discs subjected to Noc1 depletion, p53E may be selectively induced to engage stress-specific or compensatory mechanisms that modulate the nucleolar stress response in a manner that differs across tissues and developmental stages. p53E could influence the relative abundance and composition of p53 oligomeric complexes, thereby fine-tuning p53 signaling output.

In this context, p53E may alter the balance of p53A-p53B-containing complexes. Depending on the cellular milieu, p53A can affect the formation of homo- or hetero-oligomers, ultimately shaping transcriptional responses and cell fate decisions. Consistent with this model, shorter p53 isoforms have been shown to repress the activity of longer isoforms through heterotetramer formation, a conserved mechanism of p53 isoform regulation (Anbarasan and Bourdon 2019). Supporting isoform-specific regulation in vivo, Chakravarti et al. (Chakravarti et al. 2022) used isoform-specific mutants with the *hid-GFP* reporter in the germline, demonstrating differential transcription outputs among p53 isoforms. Together, these studies highlight the importance of heterotetramer formation among p53 isoforms (Zhang et al. 2014) and suggest a potential role for post-translational modifications in modulating isoform-specific functions. Further investigation will be required to determine how these regulatory mechanisms integrate under distinct cellular stress conditions, including nucleolar stress.

Post-translational modifications (PTMs) also contribute to isoform-specific regulation of p53. Indeed, ubiquitination, phosphorylation, sumoylation, and acetylation may differ among p53 isoforms due to variations in their amino acid sequences. Lok/Chk2-dependent phosphorylation at serine 4 activates DNA repair and apoptotic responses (Brodsky et al. 2004; Peters et al. 2002). The E3 ligase dSyx5 acts as a negative regulator of p53 protein levels, while the role of the deacetylase dSir2 remains unclear (Bauer and Helfand 2009). Sumoylation at lysines 26 and 302 is essential for its full transcriptional and apoptotic activity (Mauri et al. 2008).

Moreover, our findings also highlight key mechanistic differences in p53 regulation between flies and humans. In vertebrates, nucleolar stress primarily activates p53 post-translationally through protein stabilization, mediated by ribosomal proteins that inhibit the E3 ubiquitin ligase MDM2 (Lindstrom et al. 2022). In contrast, *Drosophila* lacks a clear MDM2 homolog with E3 ligase activity, suggesting an alternative regulatory mechanism. The gene *companion of reaper* (*Corp*) has been proposed as a structural and partial functional analog of mammalian MDM2. Corp binds to p53 and negatively regulates its levels through a feedback loop, similar to MDM2; however, it lacks an E3 ligase domain and thus cannot mediate canonical p53 degradation (Chakraborty et al. 2015). These distinctions point to a divergence in the evolution of p53 regulatory circuits, where transcriptional control may play a more prominent role in flies. This difference emphasizes the evolutionary divergence in p53 regulation and raises important questions about whether isoform-specific transcriptional control might also serve as a stress adaptation mechanism in vertebrates, including humans.

Further supporting the evolutionary conservation of nucleolar stress responses, our previous data show that downregulation of the human homolog of Noc1, called *CEBPZ* (Rambaldelli et al. 2025), also activates p53 both transcriptionally and at the protein level, consistent with our findings in *Drosophila* (Destefanis et al. 2022). Noc1 functions as part of a conserved nucleolar heterodimeric complex with Noc2 and Noc3, which is essential for pre-rRNA processing and 60S ribosomal subunit assembly, as demonstrated in both yeast and *Drosophila* (Milkereit et al. 2001; Destefanis et al. 2022). Our finding that Noc1 depletion triggers p53 upregulation in both flies and humans highlights the evolutionary conservation of nucleolar stress responses. This conserved activation highlights the broader relevance of nucleolar stress-induced p53 signaling and underscores the value of isoform-specific tools in elucidating the underlying regulatory mechanisms.

In conclusion, our analysis, enabled by isoform-specific primers and p53 mutant lines, reveals previously unrecognized residual activity and compensatory regulation, underscoring the intricate and multifaceted control of p53 in *Drosophila*. Our results indicate that p53E may participate in the apoptotic response triggered by nucleolar stress, suggesting it may have an isoform-specific role in this process. Furthermore, the observation that γ-H2AV phosphorylation reflects caspase-dependent apoptosis rather than direct genomic lesions indicates activation of a p53 stress response distinct from canonical genotoxic pathways. Collectively, these findings suggest that p53E is a component of a novel nucleolar stress–responsive mechanism that influences cellular adaptation to defects in ribosome biogenesis.

While our findings are consistent with potential transcriptional regulation of stress responses, further studies will be required to determine the relative contributions of transcriptional and post-translational mechanisms in maintaining cellular homeostasis.

## Data Availability Statement

All data are available in the published manuscript and its online Supplementary Files: Supplementary Figure 1, which reports the sequences of all the primers published for p53 isoforms, along with their corresponding recognition region in the *p53* locus; Supplementary Figure 2, which includes the melting curve of the primer sets used; and Supplementary Figure 3. Sequences showing the mutation present in the *p53^A2.3^*, *p53^B41.5^* mutant lines.

## Acknowledgments

We thank the Bloomington Stock Center (NIH P40OD018537). Department CIBIO Core Facilities is supported by the European Regional Development Fund (FESR) 2021–2027. The input from members of the COST Action CA21154 TRANSLACORE, supported by COST (European Cooperation in Science and Technology and by the Dipartimento di Eccellenza 2023-2027, Legge 232/2016 project n 40613, funded by the MUR. This document includes language and clarity improvements made using ChatGPT (OpenAI).

## Author contributions

AVC performed the experiments, conceived the study, and contributed to the writing of the manuscript. SB contributed to the final version of the manuscript. VM performed the experiments and analyzed the data. PB participated in the design, coordinated and supervised the team, and contributed to the writing and editing of the final manuscript. All authors have read and agreed to the published version of the manuscript.

## Bibliography

Anbarasan, T., and J.C. Bourdon, 2019 The Emerging Landscape of p53 Isoforms in Physiology, Cancer and Degenerative Diseases. Int J Mol Sci 20 (24).

Barrio, L., A. Dekanty, and M. Milan, 2014 MicroRNA-mediated regulation of Dp53 in the Drosophila fat body contributes to metabolic adaptation to nutrient deprivation. Cell Rep 8 (2):528–541.

Bauer, J.H., and S.L. Helfand, 2009 Sir2 and longevity: the p53 connection. Cell Cycle 8 (12):1821.

Boley, N., M.H. Stoiber, B.W. Booth, K.H. Wan, R.A. Hoskins et al., 2014 Genome-guided transcript assembly by integrative analysis of RNA sequence data. Nat Biotechnol 32 (4):341–346.

Boukoura, S., and D.H. Larsen, 2024 Nucleolar organization and ribosomal DNA stability in response to DNA damage. Curr Opin Cell Biol 89:102380.

Bourdon, J.C., K. Fernandes, F. Murray-Zmijewski, G. Liu, A. Diot et al., 2005 p53 isoforms can regulate p53 transcriptional activity. Genes Dev 19 (18):2122–2137.

Brand, A.H., and N. Perrimon, 1993 Targeted gene expression as a means of altering cell fates and generating dominant phenotypes. Development 118 (2):401–415.

Brodsky, M.H., W. Nordstrom, G. Tsang, E. Kwan, G.M. Rubin et al., 2000 Drosophila p53 binds a damage response element at the reaper locus. Cell 101 (1):103–113.

Brodsky, M.H., B.T. Weinert, G. Tsang, Y.S. Rong, N.M. McGinnis et al., 2004 Drosophila melanogaster MNK/Chk2 and p53 regulate multiple DNA repair and apoptotic pathways following DNA damage. Mol Cell Biol 24 (3):1219–1231.

Chakraborty, R., Y. Li, L. Zhou, and K.G. Golic, 2015 Corp Regulates P53 in Drosophila melanogaster via a Negative Feedback Loop. PLoS Genet 11 (7):e1005400.

Chakravarti, A., H.N. Thirimanne, S. Brown, and B.R. Calvi, 2022 Drosophila p53 isoforms have overlapping and distinct functions in germline genome integrity and oocyte quality control. Elife 11.

de la Cova, C., N. Senoo-Matsuda, M. Ziosi, D.C. Wu, P. Bellosta et al., 2014 Supercompetitor Status of Drosophila Myc Cells Requires p53 as a Fitness Sensor to Reprogram Metabolism and Promote Viability. Cell Metab.

Destefanis, F., V. Manara, S. Santarelli, S. Zola, M. Brambilla et al., 2022 Reduction of nucleolar NOC1 leads to the accumulation of pre-rRNAs and induces Xrp1, affecting growth and resulting in cell competition. J Cell Sci 135 (23).

Germani, F., C. Bergantinos, and L.A. Johnston, 2018 Mosaic Analysis in Drosophila. Genetics 208 (2):473–490.

Ingaramo, M.C., J.A. Sanchez, and A. Dekanty, 2018 Regulation and function of p53: A perspective from Drosophila studies. Mech Dev 154:82–90.

Lake, C.M., J.K. Holsclaw, S.P. Bellendir, J. Sekelsky, and R.S. Hawley, 2013 The development of a monoclonal antibody recognizing the Drosophila melanogaster phosphorylated histone H2A variant (gamma-H2AV). G3 (Bethesda) 3 (9):1539–1543.

Lindstrom, M.S., J. Bartek, and A. Maya-Mendoza, 2022 p53 at the crossroad of DNA replication and ribosome biogenesis stress pathways. Cell Death Differ 29 (5):972–982.

Mauri, F., L.M. McNamee, A. Lunardi, F. Chiacchiera, G. Del Sal et al., 2008 Modification of Drosophila p53 by SUMO modulates its transactivation and pro-apoptotic functions. J Biol Chem 283 (30):20848–20856.

Mesquita, D., A. Dekanty, and M. Milan, 2010 A dp53-dependent mechanism involved in coordinating tissue growth in Drosophila. PLoS Biol 8 (12):e1000566.

Milkereit, P., O. Gadal, A. Podtelejnikov, S. Trumtel, N. Gas et al., 2001 Maturation and intranuclear transport of pre-ribosomes requires Noc proteins. Cell 105 (4):499–509.

Ollmann, M., L.M. Young, C.J. Di Como, F. Karim, M. Belvin et al., 2000 Drosophila p53 is a structural and functional homolog of the tumor suppressor p53. Cell 101 (1):91–101.

Pederzolli, M., E. Barion, A. Valerio, A.V. Cuda, V. Manara et al., 2025 Optimized protocol for single-cell isolation and alkaline comet assay to detect DNA damage in cells of Drosophila wing imaginal discs. STAR Protoc 6 (1):103590.

Peters, M., C. DeLuca, A. Hirao, V. Stambolic, J. Potter et al., 2002 Chk2 regulates irradiation-induced, p53-mediated apoptosis in Drosophila. Proc Natl Acad Sci U S A 99 (17):11305–11310.

Rambaldelli, G., V. Manara, A. Vutera Cuda, G. Bertalot, M. Penzo et al., 2025 From Flies to Humans: Conserved Roles of CEBPZ, NOC2L, and NOC3L in rRNA Processing and Tumorigenesis. bioRxiv.

Robin, M., A.R. Issa, C.C. Santos, F. Napoletano, C. Petitgas et al., 2019 Drosophila p53 integrates the antagonism between autophagy and apoptosis in response to stress. Autophagy 15 (5):771–784.

Rong, Y.S., S.W. Titen, H.B. Xie, M.M. Golic, M. Bastiani et al., 2002 Targeted mutagenesis by homologous recombination in D. melanogaster. Genes Dev 16 (12):1568–1581.

Wells, B.S., and L.A. Johnston, 2012 Maintenance of imaginal disc plasticity and regenerative potential in Drosophila by p53. Dev Biol 361 (2):263–276.

Wylie, A., A.E. Jones, S. Das, W.J. Lu, and J.M. Abrams, 2022 Distinct p53 isoforms code for opposing transcriptional outcomes. Dev Cell 57 (15):1833–1846 e1836.

Zhang, B., S. Mehrotra, W.L. Ng, and B.R. Calvi, 2014 Low levels of p53 protein and chromatin silencing of p53 target genes repress apoptosis in Drosophila endocycling cells. PLoS Genet 10 (9):e1004581.

Zhang, B., M. Rotelli, M. Dixon, and B.R. Calvi, 2015 The function of Drosophila p53 isoforms in apoptosis. Cell Death Differ 22 (12):2058–2067.

