## Supplementary Fig 1 for "Noc1 Downregulation Induces Nucleolar Stress and Upregulates p53 Isoforms, with a Robust Increase of the Truncated p53E Isoform in Drosophila Wing Discs"

Supplementary Figure 1

A

| Robin et al 2019 | | |
|---|---|---|
| Target | Forward | Reverse |
| p53A | CACAGCCAATGTCGTGGCAC | GGCCATGGGTTCCGTGGTCA |
| p53B | GGACACAAATCGCAACTGCT | GGCCATGGGTTCCGTGGTCA |

| Chakravarti et al 2022 | | |
|---|---|---|
| Target | Forward | Reverse |
| p53A | CCAACAAGATCGCTTGATCAGATA | GGCCATGGGTTCCGTGGTCA |
| p53B | GAGTCAGCAGTTCGGGTCTC | GGCCATGGGTTCCGTGGTCA |

| Wylie et al 2022 | | |
|---|---|---|
| Target | Forward | Reverse |
| p53A | GGTGGCCACTACGATTCTG | GGCTATATCTGATCAAGCGATCT |
| p53B | GAGGCAACAACACGAACAAC | TGGAGTCATCCTCGGAATCA |
| p53E | CTATCAGCTCTATGAGCGCAA | GCAATAACCACCGATGTTGTG |

| Zhang et al 2014 | | |
|---|---|---|
| Target | Forward | Reverse |
| p53A | CCAACAAGATCGCTTGATCAGATA | CCACGACATTGGCTGTGATA |
| p53B | AACATGATGCAGTTCACGAACAA | TGTTATTGCCATCGGAATTATTG |
| p53 total | CCAAGCTAGAGAATCACAACATCG | TCGAGTACATCCAAAGAGACTTGG |

| AVC | | |
|---|---|---|
| Target | Forward | Reverse |
| p53A/C | TATATCACAGCCAATGTCGTG | CCTCCGTGGAGTCATCC |
| p53B | CCACAAGGGCACTGATTC | ATTCCGATCCCGATACCTC |
| p53C | GTGAGCAAATTCAAAACACGC | CCATTCGCCACGCAG |
| p53E | GAGCGCAAACAATCCGT | TGTGATTCTCTAGCTTGGGC |

B

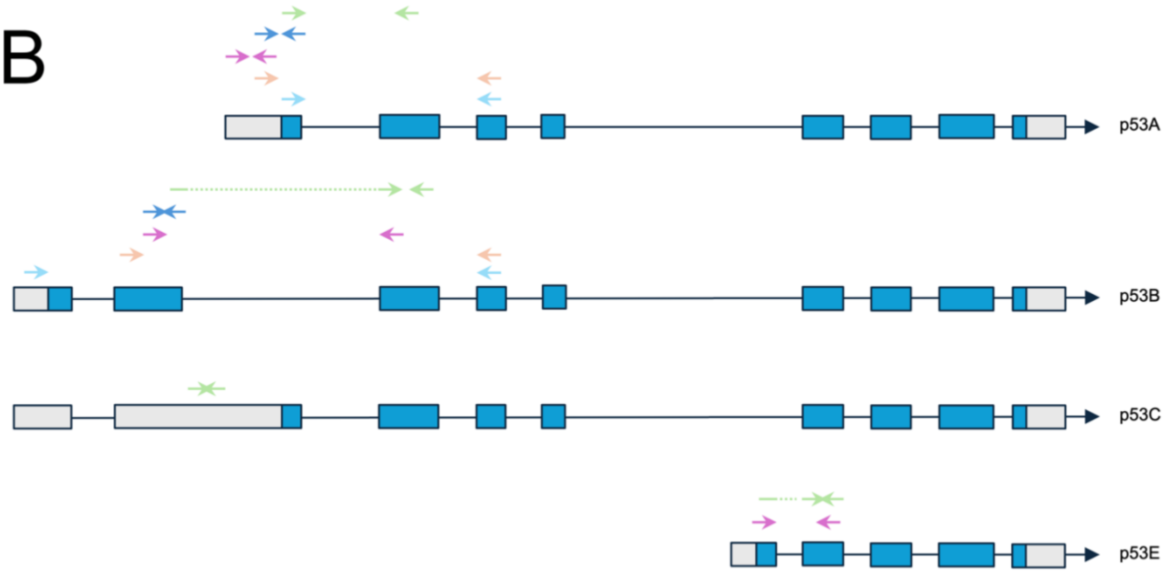


**A.** Table representing the sequences of all primers for p53. **B.** Schematic representation of p53 isoforms with the position of binding of the relative set of primers from Table A

(Robin et al. 2019; Chakravarti et al. 2022; Wylie et al. 2022; Zhang et al. 2014; Vutera Cuda et al. 2025)

Chakravarti, A., H.N. Thirimanne, S. Brown, and B.R. Calvi, 2022 Drosophila p53 isoforms have overlapping and distinct functions in germline genome integrity and oocyte quality control. *Elife* 11.

Robin, M., A.R. Issa, C.C. Santos, F. Napoletano, C. Petitgas *et al.*, 2019 Drosophila p53 integrates the antagonism between autophagy and apoptosis in response to stress. *Autophagy* 15 (5):771–784.

Vutera Cuda, A., S. Bajaj, V. Manara, and P. Bellosta, 2025 Isoform-Specific Activation of p53B and p53C in Response to Nucleolar Stress in Drosophila wing imaginal discs. *bioRxiv*.

Wylie, A., A.E. Jones, S. Das, W.J. Lu, and J.M. Abrams, 2022 Distinct p53 isoforms code for opposing transcriptional outcomes. *Dev Cell* 57 (15):1833–1846 e1836.

Zhang, B., S. Mehrotra, W.L. Ng, and B.R. Calvi, 2014 Low levels of p53 protein and chromatin silencing of p53 target genes repress apoptosis in Drosophila endocycling cells. *PLoS Genet* 10 (9):e1004581.
