## Supplementary Fig 2 for "Noc1 Downregulation Induces Nucleolar Stress and Upregulates p53 Isoforms, with a Robust Increase of the Truncated p53E Isoform in Drosophila Wing Discs"

Supplementary Figure 2

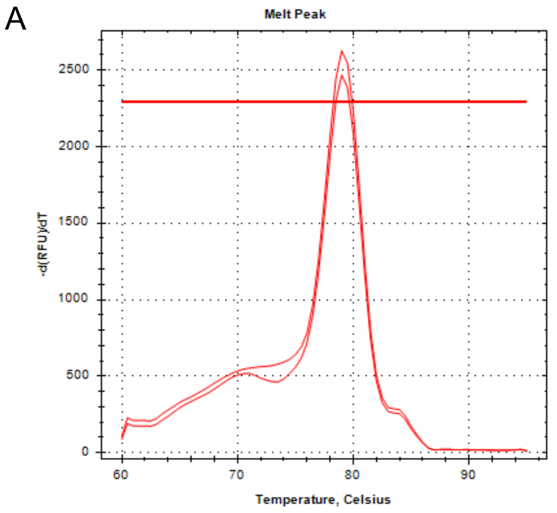


p53 pan-isoforms

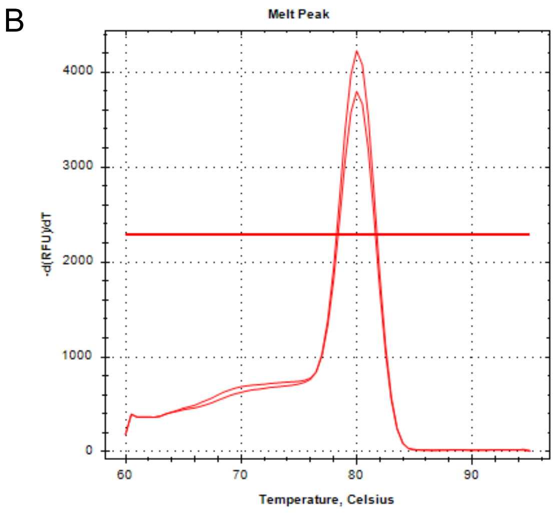


p53A/C

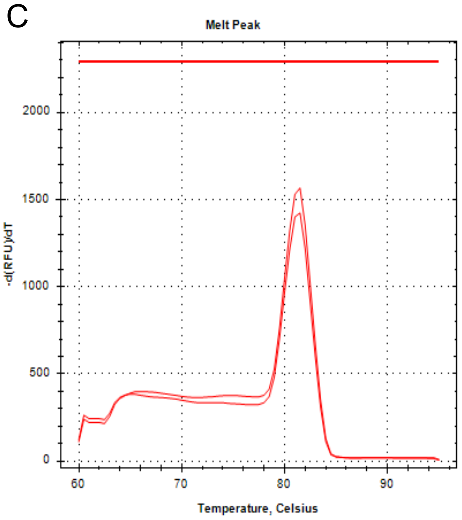


p53B

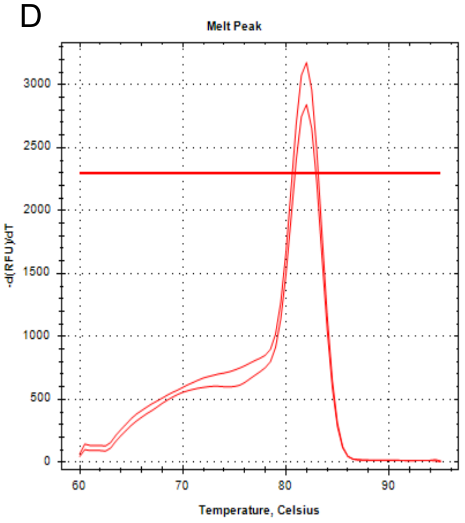


p53C

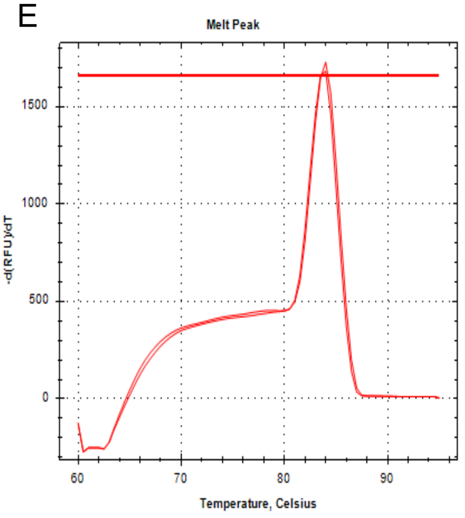


p53E

**Supplementary Figure 2. Melting curves of the p53 isoform-specific primers designed in this work.**

(A-E) Melting curves of the indicated primers obtained after qPCR on mRNA extracted from wing imaginal discs from third instar larvae from *rotund-Gal4 with UAS-Lac-z-RNAi* crosses in the $w^{1118}$ background. For every pair of primers, only one peak is present. qPCR was done with Bio-Rad CFX96, and melting curves were obtained using Bio-Rad CFX Manager software.
