## Supplementary Fig 3 for "Noc1 Downregulation Induces Nucleolar Stress and Upregulates p53 Isoforms, with a Robust Increase of the Truncated p53E Isoform in Drosophila Wing Discs"

### Supplementary Figure 3

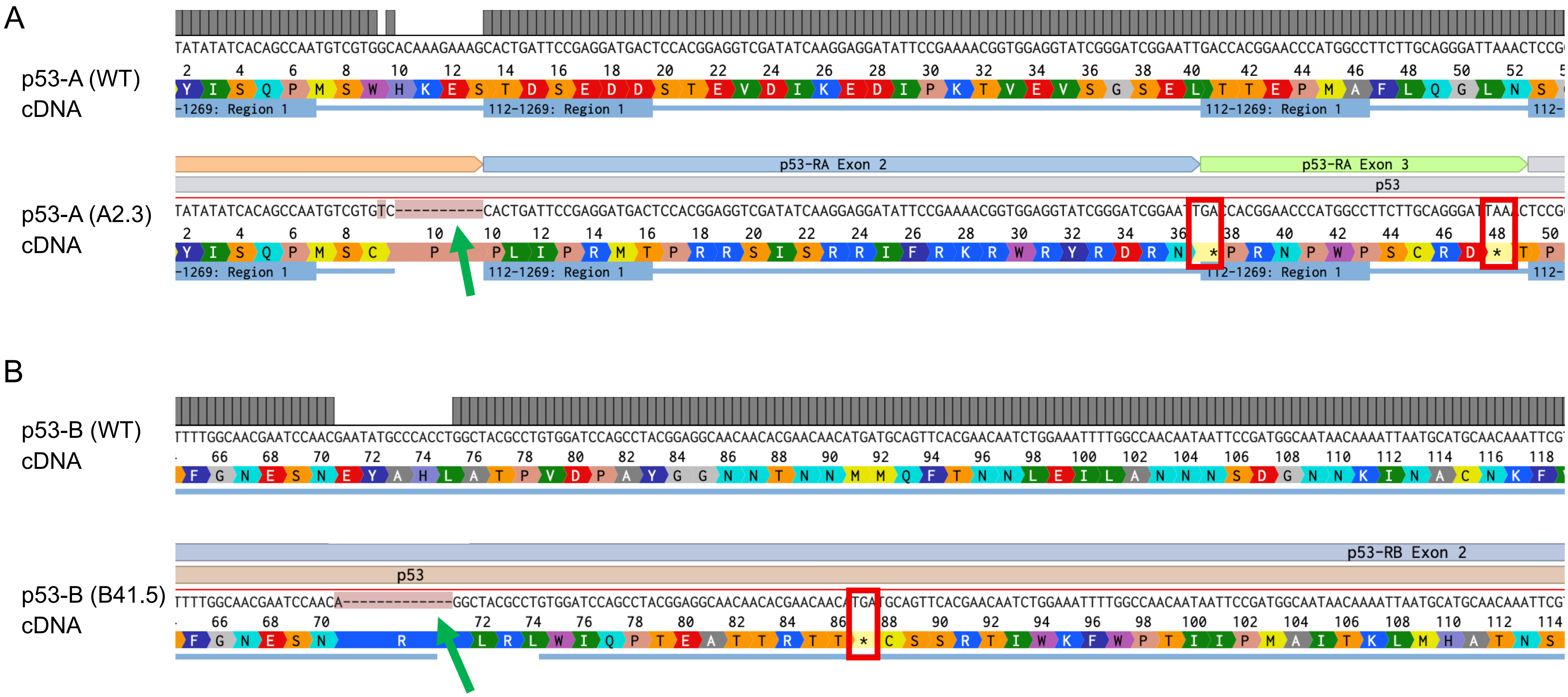


**Figure 3 (A)** Sequence alignment of the wild-type p53A region and the mutated region in the p53A (A2.3) line, showing below the translated amino acid sequence. **(B)** Sequence alignment of the wild-type p53B region and the mutant p53B (B41.5) line, showing below the translated amino acid sequence. The green arrows indicate the bases deleted in the cDNA mutant alleles. The red rectangles highlight the premature stop codons resulting from the deletion in the cDNA sequences. Alignment done in Benchling.com online software using the MAFFT algorithm with default parameters.
